# Proximal arm non-use optimises movement when the shoulder is weak: consequences for stroke patients

**DOI:** 10.1101/2020.10.26.352609

**Authors:** Germain Faity, Denis Mottet, Simon Pla, Jérôme Froger

## Abstract

Most stroke patients do not use their paretic limb whereas they are able to. The Constraint-Induced Movement Therapy (CIMT) is effective to reverse this non-use behaviour in some patients but is inapplicable or unsuccessful on others. Here, we investigate how much non-use could come from shoulder weakness instead of the behavioural conditioning treated by the CIMT. We asked 26 healthy participants to reach a target while holding a dumbbell. We found that 18/26 participants exhibit proximal arm non-use when loaded and that non-use reduces shoulder torque of final posture. We either found that non-use improves accuracy in a high gravity field. Following optimal control policy, we explain how the non-use could be an adaptative solution when the shoulder is weak. Our results show the need to include muscular strength into cost function used to model human movement. The framework presented here suggests that psychological non-use could be treated effectively with CIMT, while physiological non-use, resulting from shoulder weakness, might respond better to anti-gravity muscles strengthening.

## 1. Introduction

Stroke is one of the leading causes of upper-limb disability. In 2002, Sterr *et al*. found that only 22% of people with stroke would open a window with their paretic limb whereas 90% could do it with a sufficient quality of movement. When patients no longer use their paretic arm in favour of the less affected arm, stimulations of the remaining sensorimotor areas decrease, leading to a secondary repression of the paretic arm sensorimotor areas (Taub et al., 2002). As a result, the more the patient neglects its paretic arm, the more likely he is to lose his motor ability permanently, as summarised by the saying “use it or lose it” (Hidaka et al., 2012). This mechanism is described by Taub as the learned non-use phenomenon.

Constraint-induced movement therapy (CIMT) have been developed to counteract the learned non-use effect (Taub et al., 2002). During 14 days, less affected arm is sustained in a sling, which forces the patient to re-use its paretic arm. This intensive use mixed with the non-use of the less affected arm reverses the learned non-use phenomenon at a behavioural and at a cortical level (Liepert et al., 2000). Despite its effectiveness compared to conventional therapy (Taub et al., 2006; Wolf et al., 2006), CIMT suffers from several limits. First, because the principle is to ask people who struggle to perform their daily-life activities to no longer use their less affected arm, a residual ability to use the paretic arm is mandatory to apply CIMT. This difficulty limits inclusions (6.5% of eligibility) and might causes poor adhesion to the program (Fabbrini et al., 2014; Kwakkel et al., 2007). Second, as with all treatments, some patients do not respond, potentially masking an improved efficacy in responding patients (Fritz et al., 2012). Third, CIMT improves the upper limb function but not always the performance nor the ability (Corbetta et al., 2015). Some researchers have even reported an increase in proximal compensation at the end of the CIMT: because the less affected arm is no longer available, patients use available compensatory degrees of freedom (DoF) like trunk anterior flexion or shoulder abduction (Kitago et al., 2013; Massie et al., 2009). If it turns out that the compensation is unnecessary, this shift could lead to proximal arm non-use (PANU) as in more than half hemiparetic patients (Bakhti et al., 2017). Faced with these observations, Fritz (2012) recommends targeting well respondent patients to better allocate money and time. Other patients might instead better respond to other treatments.

Paresis, especially shoulder and elbow weakness, is a major factor of motor trouble post-stroke (Harris & Eng, 2007; Raghavan, 2015). In addition, McCrea et al. (2005) showed that shoulder flexors are often maximally activated during a reach to grasp movement in hemiparetic patients. As a result, and accordingly to optimal control logic, when the non-paretic hand is blocked but that the trunk is free, patients suffering from shoulder weakness might unconsciously move the trunk or abduct the shoulder to unload the shoulder flexion and thus achieve the task with minimum effort.

Why do more than half of stroke patients exhibit proximal compensations when they can perform the task without? Here, we hypothesise that the lack of force of the antigravity muscles of the upper limb explains the non-use of the proximal joints in people with stroke. Because the optimal control logic holds for patients and healthy people as well, we investigate whether a greater weight-over-force ratio of the arm in healthy participants could increase proximal arm non-use (PANU).

## 2. METHODS

### 2.1. Participants

Twenty-six healthy participants (12 males, age 21 ± 3 years, 3 left-handed, height 1.73 ± 0.09 m, weight 66.92 ± 9.29 kg, shoulder flexion maximum voluntary contraction (MVC) 10.33 ± 3.12 kg) participated in this study. Participants were excluded if they had shoulder pain or any other problem that could affect the reaching task. Written informed consent was obtained from all participants before their inclusion. This study was performed in accordance with the 1964 Declaration of Helsinki. Institutional Review Board of EuroMov from Montpellier University (IRB-EM 1901C) approved the study.

### 2.2. Experimental setup

Hands and trunk movements were recorded with 8 infrared optical cameras from the Vicon Motion Capture System (Vantage V5, lens 8.5 mm, Oxford Metrics, UK). The sampling frequency was set at 100 Hz. VICON Nexus 2 software was used to save time series of each marker. Experimenter placed markers on the target and the manubrium, and for each body side on the 1^st^ metacarpal, the lateral epicondyle of the humerus, the acromion process, and the iliospinal. For each side, we corrected iliospinal marker position before data analysis to correspond to the anatomical centre of hips joints.

### 2.3. Procedure

Twenty-six healthy participants performed a seated reaching task with a high weight-over-force ratio of the arm. To induce a high ratio, participants did the task while holding a dumbbell. Hand assessed was randomly selected (12 left, 14 right). The weight of the dumbbell was adjusted to 85% of the MVC. As a consequence, the shoulder torque needed to hold the arm straight next to the target was 13.0% (3.7) of the maximum in control condition against 75.0% (5.5) in the loaded condition. MVC was previously determined in a seated position. Participants had to hold the dumbbell in front of them, arm extended and pull up as much as possible for 3 seconds. The dumbbell was linked with a static rope to a force sensor located in the ground. MVC was set as the sum of the force detected by the force sensor and the weight of the dumbbell. Participants had to stay in contact with the backrest during the entire contraction. Any help of the contralateral arm was prohibited. Experimenter encouraged verbally the participant. We retained the maximum score over 3 trials separated with a rest time of 1 minute.

The main task was to reach a target located in front of the participant, at the length of the arm, like described in a previous study (Bakhti *et al*., 2017). The starting position was seated, feet on the ground, back in contact with the chair and forearm on the armrest. To assess the spontaneous arm use, participants had to reach the target in a natural pace, wait for 1 second and come back to the starting position. To assess the maximal arm use, participants had to perform the reaching task while minimising trunk compensation. Experimenter manually applied a light proprioceptive feedback on the participant’s shoulders in the maximal condition as a reminder to avoid trunk movement. Participants performed 5 trials in the control condition, then 5 trials in the loaded condition.

### 2.4. Data analysis

Data analysis was processed with SciLab 6.0.2. A low-pass Butterworth digital filter of 2nd order of 5 Hz and a cut-off frequency of 30Hz were applied on the signal. First, we computed the beginning and end of each reaching movement from the hand maker time series. The beginning of the movement was when the Euclidean velocity became positive and remained positive until the maximum velocity. The end of the movement was when the Euclidean distance to the target reached its minimum. Second, we computed Proximal Arm Use (PAU) for each trial as (hand displacement - trunk displacement) / hand displacement. We finally computed PANU as Maximal PAU minus Spontaneous PAU (Bakhti *et al*., 2017). Median PANU trial was retained for data analysis. Accuracy was assessed through end point error, computed as the Euclidean distance between the active hand and the target at the end of the reaching movement. Efficiency was assessed through movement time and path length ratio (Schwarz *et al*., 2019). Joints involvement was assessed through angular rotations, computed as the difference of angular position between final posture and starting posture. Elbow and shoulder torque were computed as 3D static torque of each joint at final posture. In order to compute static torque, positions of centre of mass and absolute weight of upper limbs were approximated from height and weight of participants following De Leva’s equations (1996).

### 2.5. Statistical analysis

The statistical analysis was conducted using R 3.6.1 with rstatix and stats packages. Density plots and quantiles/quantiles plots revealed a violation of the normality assumption inside data groups. Therefore, all the hypotheses were studied in a non-parametric way. PANU was studied with the Wilcoxon signed rank test for the side factor (within-subject, 2 modalities: control VS loaded). Elbow and shoulder torque, angles, movement time, end point error and path length ratio were studied with the Wilcoxon signed rank test for the weight factor (within-subject, 2 modalities: control VS loaded) and the arm-use factor (within-subject, 2 modalities: spontaneous use VS maximal use). Because of the multiple comparisons, p-values were corrected by the method of Benjamini & Hochberg (1995). The significance level was set at .05 in all analysis. The effect size r was calculated as Z statistic divided by square root of the number of observation (N): r = Z / √N (R. Rosenthal et al., 1994). Data in the text are presented as median (interquartile range).

## 3. RESULTS

### 3.1. Increasing the weight-over-force ratio of the arm induces proximal arm non-use in healthy participants

A Wilcoxon Signed-Ranks Test indicated that the PANU scores in the loaded condition (11.91%, IQR = 14.23) were greater than in the control condition (1.14%, IQR = 0.75), W = 0, p <.001, r = .87 (figure 2). This result indicates that a weight-over-force ratio of 75.0% (5.5) (compared to 13.0%, IQR = 3.7) induces PANU in 69% of healthy participants.

#### Effect of the increase of weight-over-force ratio on the reaching coordination

A detailed analysis shows that the spontaneous reach in control condition exhibited an elbow extension of 34.04° (15.74), a shoulder flexion of 39.73° (9.56) and a shoulder adduction of 21.05° (7.20). Trunk was also involved with an anterior trunk flexion of 0.61° (0.64) and a trunk rotation of 6.06° (3.00) (figure 3).

When participants performed the same task with an increased weight-over-force ratio, they produced 57% less elbow extension (W = 343, p <.001, r = .83), 18% less shoulder flexion (W = 304, p < .01, r = .64) and 65% less shoulder abduction (W = 19, p < .001, r = .78). Loaded participants compensated using 710% more anterior trunk flexion (W = 11, p < .001, r = .82) and 188% more trunk rotation (W = 38, p < 0.001, r = .68).

#### Effect of trunk restriction on the reaching coordination

Minimising trunk use during loaded movement has the effect to narrow the gap between control and loaded coordination. There was an expected decrease of anterior trunk flexion (−76%, W = 0, p < .001, r = .87) and rotation (−38%, W = 38, p < .001, r = .68). In addition, trunk restraint was associated with an increased elbow extension (+111%, W = 331, p < .001, r = .77) and a non-significant increased shoulder flexion (+15%, W = 261, p = .058, r = .43). On the other hand, shoulder adduction was not modified (+34%, W = 191, p = .708, r = .08). Summing up, participants with a high weight-over-force ratio decrease elbow extension and increase trunk use. In addition, they kept their shoulder abducted instead of the adduction seen in the control condition.

Less expected, minimising trunk use during control movement had also the significant effect to diminish trunk flexion (−69%, W = 15, p < .001, r = .80) and trunk rotation (−42%, W = 6, p < .001, r = .84), at the expense of elbow extension (+29%, W = 349, p < .001, r = .86) and shoulder adduction (+3%, W = 85, p < .05, r = .45). Nevertheless, shoulder flexion increase did not reach significance (+7%, W = 250, p = .059, r = .37).

### 3.2. High weight-over-force ratio induces a slower and less accurate reach

The increase of weight-force-ratio of the arm had also made the movement slower and less efficient in both spontaneous and maximum conditions. However, the restriction of the trunk had no effect on the time of movement and efficiency.

Spontaneous movements were slower in the loaded condition (2.29 s, IQR = 0.64) compared to the control condition (1.68 s, IQR = 0.82) (W = 57, p < .01, r = .59). In addition, maximal movements were slower in the loaded condition (2.21 s, IQR = 0.84) compared to the control condition (1.68 s, IQR = 0.54) (W = 71, p < .05, r = .49). We found no difference of time movement between spontaneous and maximal conditions (control: W = 148.5, p = .72, r = .07; loaded: W = 146.5, p = .72, r = .09).

Path length ratio were greater in the loaded condition than in the control condition for both spontaneous and maximal movements (control spontaneous: 1.08 (0.07); loaded spontaneous: 1.14 (0.12); W = 23, p < .001, r = .69; control maximal: 1.08 (0.08); loaded maximal: 1.18 (0.12); W = 11, p < .001, r = .79). Again, we found no difference between spontaneous and maximal conditions (control: W = 140, p = .40, r = .16, loaded: W = 208.5, p = .19, r = .33).

On the other hand, trunk restraint diminished accuracy when reaching with a high weight-over-force ratio (end point error spontaneous: 29 mm (10), maximal: 34 mm (22), W = 270, p < .05, r = .47). Surprisingly, participants in control condition were more accurate during trunk restraint than during spontaneous movement (end point error spontaneous: 31 mm (6); maximal: 29 mm (6); W = 54.5, p < .05, r = .54). Finally, arm weight increase did not affect accuracy (spontaneous: W = 197.5, p = .18, r = .26; maximal: W = 90.5, p = .06, r = .42).

### 3.3. Non-use coordination reduces shoulder torque of final posture

Trunk restraint increased shoulder torque of final posture by 10% in control (spontaneous: 7.53 N.m (2.21); maximal: 8.25 N.m (2.62); W = 351, p < .001, r = .87) and by 16% in loaded condition (spontaneous: 36.75 N.m (14.33); maximal: 42.81 N.m (21.85); W = 351, p < .001, r = .87). In other words, trunk restraint increased the mechanical cost needed at the shoulder to maintain the arm in its final position. Given the increased arm weight in the loaded condition, we made no comparison between control and loaded conditions.

We found no effect of trunk restraint on elbow torque both in control (spontaneous: 2.36 N.m (0.89); maximal: 2.38 N.m (0.85); W = 227.5, p = .055, r = .45) and loaded condition (spontaneous: 24.09 N.m (13.83); maximal: 23.23 N.m (14.01); W = 194, p = .653, r = .09).

## 4. DISCUSSION

In this study, we show that increasing the weight-over-force ratio of the arm induces proximal arm non-use (PANU) in 69% of healthy participants. We either found that trunk compensation reduces shoulder torque and increases accuracy of the reach, without affecting movement efficiency. We explain proximal arm non-use (PANU) as an optimal coordination pattern when antigravity shoulder muscles are weak, both in healthy and stroke participants.

### 4.1. Neuromuscular weakness induces proximal arm non-use: optimal control point of view

When a seated healthy participant has to reach a target in front of him (*e.g*. a glass of water), he has to meet the constraints of the task and of his body. So, the goal of its movement is to move his hand from the armrest to the target, in a way that grabbing the glass is possible: with hand open and palm in front of the glass. But he also has to meet the constraints of his body: he cannot extend the elbow more than 180°, lift 1000 kg, or execute the movement in less than 20 ms. Its movement is capped by the intrinsic properties of its neuro-musculoskeletal system, such as articular range, force, and velocity. Thanks to the many degrees of freedom of the human body, there are still endless ways to perform the movement by meeting constraints of the task: the human body is redundant. But how is chosen the coordination finally performed, among the infinite possibilities? Optimal control logic suggests that the human system performs the coordination that minimise a certain cost (Todorov & Jordan, 2002). This cost is related to the human effort (Venture et al., 2019) but there is no consensus about its exact nature. Consequently, we will first consider it in a general way before discussing the contribution of our experiment to the optimal control framework.

Whatever the cost looked at, healthy participants exhibit a “normal way” when reaching, *i.e*. flexing and adducting the shoulder while extending the elbow (figure 1b). Little mass is moved, it costs little effort to move the arm to the target. Because the target is further than 80% of the upper limb length, participants use minor trunk anterior flexion and rotation (Robertson & Roby-Brami, 2011) (figure 3). Still in control condition, when we ask the participant to not use trunk movement, a larger elbow extension and shoulder adduction compensate for the smaller trunk flexion and rotation (figure 1a, 3). This shift is reflected by a median PANU of 1.14% (0.75) in control condition, instead of 0 in the ideal situation in which trunk would be perfectly motionless (figure 2). Moreover, the shift of work from trunk to upper limb increases shoulder torque by 10 %. This shift of coordination seems too light to induce an effect on movement efficiency, although the constraint increases accuracy of the reach. In summary, restraining the trunk during a reaching movement in healthy participants increases the upper limb work without degrading the performance.

**Figure 1.**
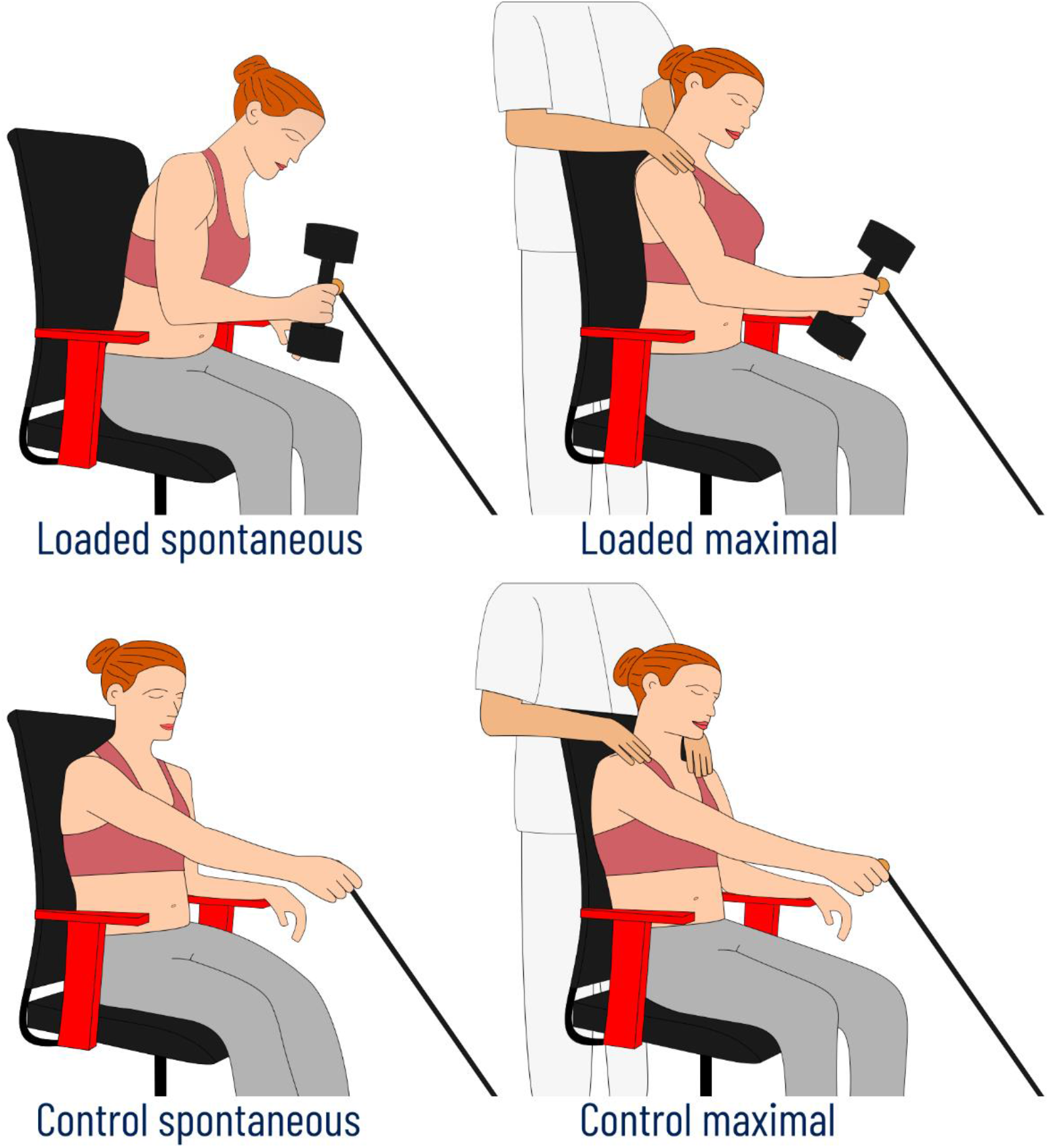
Schema of experimental conditions.

**Figure 2.**
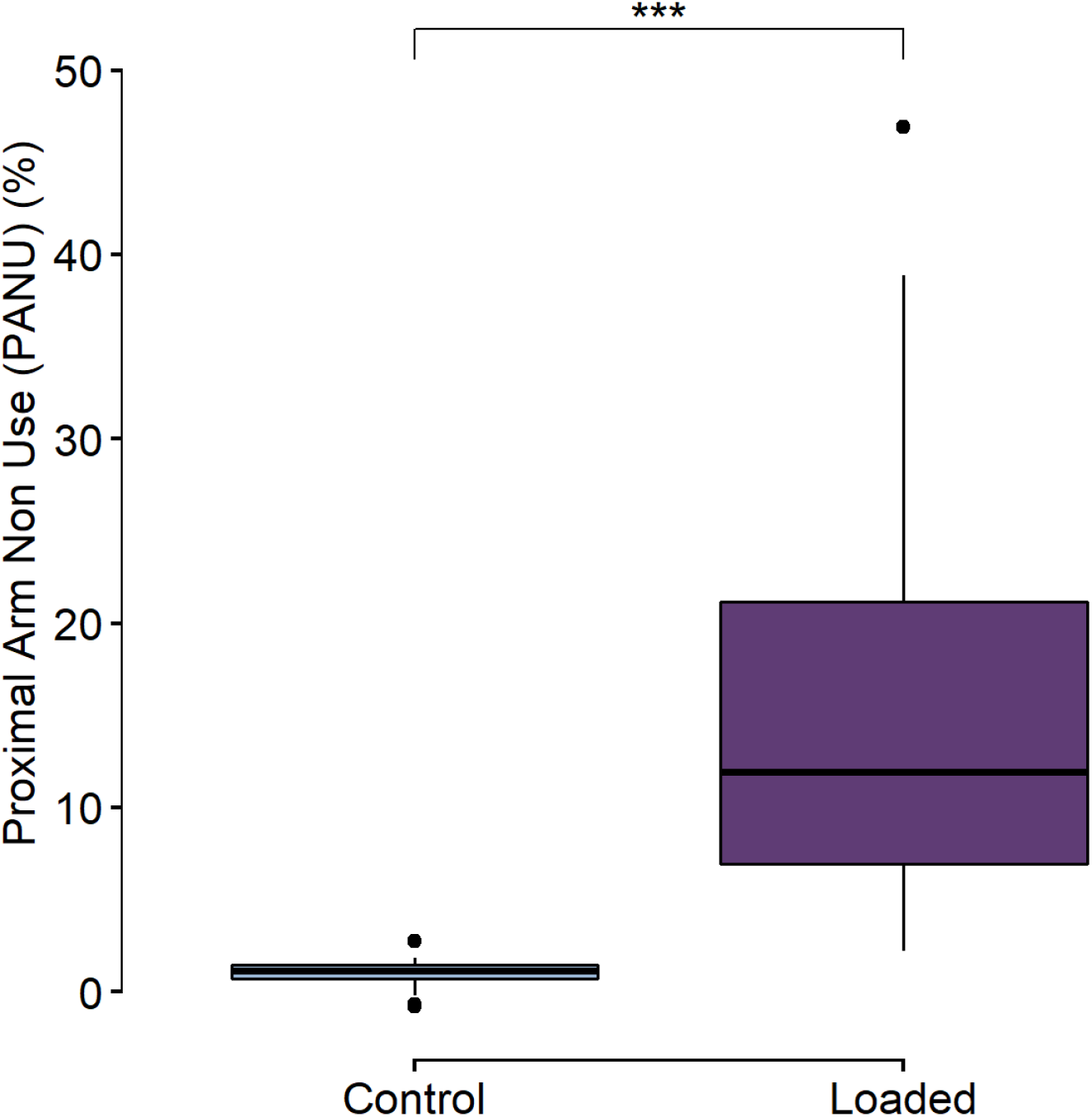
Proximal arm non-use (PANU) in control and loaded condition. Internal horizontal bar represents median, boxes extend from first to third quartile, whiskers extend from minimal to maximal values excluding outliers and white circles represent outliers.

**Figure 3.**
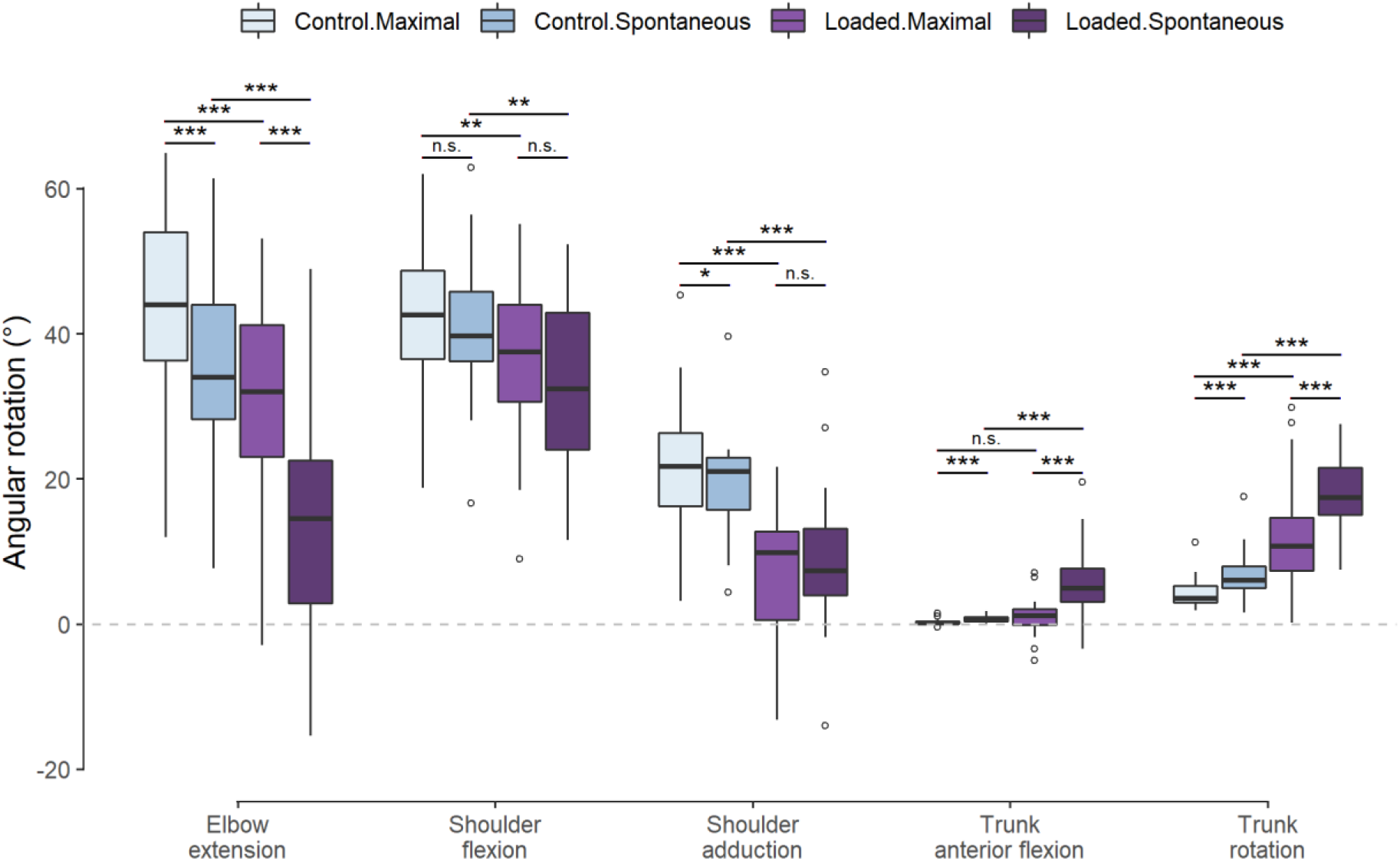
Differences of angular rotation between final and starting posture. Negative values represent inverse behaviour (for example, shoulder abduction instead of shoulder adduction during movement). Internal horizontal bar represents median, boxes extend from first to third quartile, whiskers extend from minimal to maximal values excluding outliers and white circles represent outliers.

The reduced end point error in control-maximal condition could come from confounding factors and should be replicated before any extrapolation. First, p-value is not far from non-significance (p = .013) compared to other p-values that largely exceed significance. Second, spontaneous condition is always performed before maximal condition to guarantee the integrity of the movement spontaneity. So, participants had already done 5+ trials of each hand before performing the maximal condition against only a few before the spontaneous condition. Thus, this result could be caused by a learning effect.

However, in loaded condition, the torque needed to maintain the arm straight next to the target is 75.0% (5.5) of the maximum torque producible by the participant, against only 13.0% (3.7) in control condition. Thus, when the participant lifts his arm straight in front of him, he is near to his maximum of neuromuscular activation: it is challenging to him to lift his own arm. The task is still the same, *i.e*. moving the hand from the armrest to the target, the control laws are still the same, *i.e*. minimising effort, but the intrinsic parameters of the system have changed. Despite the instruction to minimise trunk movement, the increased weight-over-force ratio of the arm induces a greater involvement of compensation pattern, such as freezing shoulder abduction and rotating the trunk to bring the shoulder - and so the hand - closer to the target (figure 1c). In addition, when the participant is loaded, his antigravity muscles are activated near the maximum. Because increasing the level of activation increases the signal-dependent noise (Jones et al., 2002), the reach becomes more noisy. As a result, the participant has to perform more corrective feedback loops during movement, which segment the path. Moreover, at each correction the velocity slow-down before increasing again. In accordance to the longer path and the slower velocity, the movement time increases.

But loaded participants take advantage of the redundancy of the human body. By using 312% more trunk anterior flexion and 63% more trunk rotation when their trunk is free, they diminish elbow extension and shoulder flexion, which has the effect to reduce the distance between hand and shoulder (figure 1d, 3). The elevation of the arm is then completed by freezing the shoulder in abduction. The reduced arm lever results in a reduced shoulder torque (−16%) and so in less effort to achieve the task. Because a reduced muscular activation minimises signal-dependent noise, the non-use behaviour improves accuracy of loaded participants by 13%. Trunk involvement in both control and loaded condition illustrates the principle of distribution of work across multiple effectors: “optimality is achieved when the system distributes work across multiple effectors, even if task goal could be achieved by the activation of a single muscle” (Diedrichsen et al., 2010). Here, in both control and loaded condition, PANU minimises effort while maximising accuracy of the reach. Proximal arm non-use is thus an adaptive way of reaching especially when gravity force becomes difficult to counteract.

### 4.2. Post-stroke movement still follows optimal control laws

In the previous section, we have demonstrated that the coordination of reaching is located on a continuum between an exclusive use of the upper limb and a shared contribution between elbow, shoulder and trunk use. The location of the coordination on this continuum is defined by the task condition (trunk restraint or trunk use) and by the shoulder strength of the participant. Thus, an “upper limb focus” involves the displacement of fewer masses and so a reduced mechanical cost whereas a “trunk focus” produces a stronger and more accurate movement. This continuum is similar to the one found in stroke patients: most affected patients exhibit larger trunk involvement than mild patients (Cirstea & Levin, 2000). In fact, upper limb paresis following a stroke induces a high weight-over-force ratio of the arm (Ada et al., 2006; Bourbonnais, Noven, 1989). As a result, stroke patients are near the maximum of neuromuscular activation during a reaching movement (McCrea et al., 2005). By compensating with the trunk, patients may successfully reduce shoulder torque and thus minimise effort while improving accuracy. Indeed, reducing shoulder torque in stroke patients with arm weight support successfully reduce shoulder neuromuscular activation (Coscia et al., 2014; Runnalls et al., 2019), and results in a larger reachable space (Dewald et al., 2001; Sukal et al., 2007). As a consequence, on a single reach, it may be more optimal to freely use all the available degrees of freedom instead of spending more attention, effort, energy to perform the task more “normally”. Thus, trunk involvement and shoulder abduction, shared by healthy and stroke participants, is not ineluctably a “bad synergy” resulting from pathological reflexes association or impaired control (Dewald et al., 1995; Subramanian et al., 2020), but rather testifies that stroke movements continue to follow optimal control laws, as claimed by Latash & Anson since 1996. Our conclusions are consistent with the energy minimization during the walk in stroke patients (Roemmich et al., 2019).

However, restraining non-use could have a positive long-term effect, such as avoiding repression of the paretic arm cortical zones, and increasing effort during reaching could be the price to pay. Indeed, the CIMT logic is to restrain the non-paretic upper limb in a sling for 14 days to force the patient to re-use its paretic arm in his everyday life. This type of therapy show a medium effect size (standard mean difference of .34) on arm motor function compared with conventional treatment (Corbetta et al., 2015), but its effectiveness might be limited to patients who already have sufficient upper limb strength.

In fact, if the patient has a near normal shoulder strength, the compensation is probably not mandatory. In this case, he might succeed in re-using paretic hand without significantly degrading his performance. According to Taub (2002), the non-use repressed may be a psychological residual from the acute phase of the stroke.

But other patients could have a high, but not maximal, weight-over-force ratio (<100%). As seen in our results, in this case, the compensation is not mandatory, but is an optimization to the patient’s condition. So, when the non-paretic hand is restrained, patients experience a movement less accurate and more effortful. As a result, patients exhibit proximal compensations, such as trunk involvement and shoulder abduction, to optimise their movement (Kitago et al., 2013; Massie et al., 2009). This strategy is effective to improve motor function of the paretic arm but can lead to proximal arm non-use. Because of its implication in the optimization of the movement, we could name this type of non-use a “physiological non-use”.

Some therapies have been developed precisely to reduce proximal compensation. As seen in our results, trunk restraint (TR) does decrease trunk compensation at the expense of elbow-shoulder recruitment during the task (Michaelsen et al., 2001; Pain et al., 2015). In addition, when they follow trunk restraint therapy, patients keep this shift of coordination at least 1 month after the program (Michaelsen et al., 2006). Similar results have been described with trunk movement-oriented biofeedback (Valdés et al., 2017). As a result, a combination of CIMT and trunk restraint is more effective in decreasing compensations than CIMT alone (Bang et al., 2018; Wu et al., 2012). Nevertheless, the compensatory use of shoulder abduction might persist when the shoulder is weak (figure 3). In addition, TR has so far failed to demonstrate any improvement of upper body limb function, smoothness and straightness in stroke movement (Wee et al., 2014). To conclude, therapies exist and are effective in reducing non-use in patients suffering from shoulder weakness, but at the price of sub-optimal movements. Unfortunately, we do not have enough data yet on comparing quality of life at long term, the only factor that really counts.

Although CIMT and TR might be effective in patients suffering from a psychological or a physiological non-use, they may be ineffective in patients that have not enough shoulder strength (ratio < 100%): we cannot remove a mandatory compensation. In this case, other treatments should be applied.

### 4.3. Using optimal control laws to improve stroke rehabilitation

If the patient exhibits the same level of non-use whenever its paretic limb is lightened, then its shoulder torque does not influence non-use: non-use is psychological. In this case CIMT should successfully increase paretic arm use and TR should decrease the proximal arm non-use without degrading upper limb performance, at least in the long term.

But if the non-use decreases when the paretic limb is lightened, then part of the non-use is physiological: the patient compensates to counteract the negative effects of the shoulder weakness. In this case, CIMT and TR would be less effective in decreasing non-use and could degrade performance. Another solution would be to increase anti-gravity shoulder muscle force. Thus, instead of forcing an atypical system to commit a normal coordination, the increased force would directly modify the system parameters into more normal values. As a consequence, the optimal coordination would become the one without compensations. This is supported by the fact that upper limb strength explains 79% of variance of arm activity after a stroke (Harris & Eng, 2007). It is therefore important to better evaluate therapies capable of improving stroke patients’ strength. For example, resistance training is promising (Ada et al., 2006; Harris & Eng, 2010) but scientific evidence suffers from a lack of standardization which prevents us from conclude on its efficacy (Gambassi et al., 2017). Therapists assessing functional strength training intervention should follow TIDieR checklist (Vliet et al., 2016). However, outcomes show a reduction in compensatory movements after upper limb progressive resistance therapy (Ellis et al., 2018; Thielman et al., 2004). Because resistance training gains are very specific, it might increase efficacy to mix resistance training with task oriented repetitive training, as adopted in the promising study of Högg *et al*. (2019). Afterwards, if residual non-use persists even after a sufficient increase in the weight-over-force ratio, then psychological non-use therapies could be applied (CIMT, TR, trunk compensation biofeedback).

Finally, if the weight-over-force ratio of the arm is too high (>100%) to induce a non-use, shoulder strength increase is mandatory to diminishes compensations. Effective therapies to increase strength should be assessed. When applying a strengthening program, therapists should therefore consider arm weight into computation of work load, especially when shoulder is very weak. For example, if the patient could lift only 80% of its arm weight, then resistance training at 70% of MVC should be done at 80% * 70% = 56% of arm weight. Training could then be done with a pulley system to lighten the arm of the patient. Another solution is to decrease the weight-over-force ratio using wearable exoskeletons (Gassert & Dietz, 2018). In certain patients, an increase of some newton-meters in the shoulder torque might promote the paretic arm use in daily life.

### 4.4. Theoretical implications for motor control

Optimal control theory is based on the minimization of one or more cost functions (Todorov, 2004). Thus, coordination is performed to satisfy the constraints of the task while minimising a cost. Objective cost functions have been first explored to minimise a mechanical cost (Nelson, 1983; Uno et al., 1989), but according to Loeb (2012), cost functions not based on subjective feelings make no sense in a neurobiological system. More human-oriented functions have then been developed, such as minimum neuromuscular effort (Guigon et al., 2007), minimum metabolic cost (Praagman et al., 2006) or maximum comfort (Cruse et al., 1990). Finally, other functions combine objective and subjective effort (Wang et al., 2016) or introduce a modulation by individual traits (Berret et al., 2018; Rigoux & Guigon, 2012). In spite of this diversity, there is currently no consensus on any cost function.

Here, we demonstrate that the modification of the end-effector mass leads to a shift of coordination. As we obtain a change in coordination similar to that observed in the stroke population, the common factor (high weight-over-force ratio of the arm) might be responsible for the change. Known cost functions should be evaluated for their ability to consider available muscle strength. Individual characteristics, such as psychological traits or previous experiences, may explain the diversity of behaviours under similar physical conditions.

## 5. CONCLUSION

Upper limb non-use, a common outcome after a stroke, may exacerbate motor deficit in the long term. Constraint induced motor therapy (CIMT) is effective to enhance paretic limb activity but have a poor rate of eligibility (6.5%). In addition, some patients exhibit an increase of proximal compensation after CIMT. Here we demonstrate that a low weight-over-force ratio of the arm can exacerbate proximal arm non-use in healthy participants. In contrast to the psychological non-use described by Taub, which is insensitive to gravity, there may be a form of physiological non-use in stroke patients. Thus, a decrease of non-use with arm support would explain the compensation shift: physiological non-use improves accuracy of the reach while minimising effort against gravity. If this type of adaptative non-use is confirmed in future work, shoulder strength-oriented therapies might be considered as an alternative to CIMT in patients exhibiting physiological non-use.

## 8. Ethics approval

This study has been performed in accordance with the 1964 Declaration of Helsinki. Institutional Review Board of EuroMov from Montpellier University (IRB-EM 1901C) approved the study.

## 9. Consent to participate

Informed consent was obtained from all individual participants included in the study.

## 10. Consent for publication

Not applicable.

## 11. Availability of data and material

The raw datasets generated during the current study are available in the Open Science Framework repository: https://osf.io/r3xcu/

## 12. Code availability

The code generated during the current study is available in the Open Science Framework repository: https://osf.io/r3xcu/

## 13. Authors’ contributions

GF designed the protocol, recorded and analysed the data, wrote the first version of the manuscript. DM and JF assisted GF in designing the protocol, analysing the data and provided guidance for improving the manuscript. SP assisted GF in setting up the protocol, recording the data and improving the manuscript. All authors accepted the latest version of the manuscript.

## Acknowledgements

Not applicable.

